# Game theoretic consideration of transgenic bacteria in the human gut microbiota as a pro-biotic prophylactic for metabolic syndrome

**DOI:** 10.1101/206227

**Authors:** Ahmed M. Ibrahim, James Smith

## Abstract

A game theoretic treatment is introduced to explore the role of nutrition of the gut bacterial microflora as a potential pro-biotic therapy. Rational design of functional foods and nutraceuticals has far reaching public health and therapeutic benefits. Understanding ecological dynamics and how phenotypic manipulations of microbe-microbe interactions of the gut microbiota can provide direct health benefit is currently a fundamental question in bioengineering and more widely in the food, diet and health industries. This work considers a hypothetical adjustment of the microbiome by introducing a transgenic bacterial species that contributes to increased exposure of omega 3 fats in the gut by converting them from the omega 6 fats, dominant in the Western diet. The ratio of the two fats circulating in blood are risk markers, indicators of metabolic syndrome and related conditions. In this work, we consider nutritional exposure to a pro-biotic, a live culture of transgenic bacteria contributing omega 3 fats from omega 6 in the diet. Maintaining a long-term co-existence between native (indigenous) and transgenic bacteria is a challenge. Game theory is the appropriate tool for handling this conflict. The long-term co-existence is guaranteed if the two strains engage in the Snowdrift game. Our game theoretic treatment provides the basis of a model mechanism for prophylactic nutritional therapy for metabolic syndrome by the transgenic bacteria, providing support of indigenous gut microbiota and additional supplementation of a pro-biotic.

## Introduction

A game theoretic treatment is introduced to explore the role of nutrition of the gut bacterial microflora as a potential pro-biotic therapy for addressing the absorption of polyunsaturated fats (PUFAs) that are dominated by omega 6 (n6) class over the omega 3 (n3) class. A high n6 / n3 ratio is an indicator for high-risk metabolic syndrome, pre-type 2 diabetes and cardiovascular disease.

Metabolic syndrome is a cluster of strongly associated diseases including systemic cardiovascular disease, hypertension, osteoarthritis, stroke [1], lipodystrophies, pre-diabetes and obesity and a wide range of cancer types [2, 3]. Long-term increased dietary exposure of n3 and the increased n3/n6 ratio of circulatory fats are considered to have protective benefits and to the colon [4, 5, 6, 7]. However, the *Western diet* is typically dominated by n6 and depleted in n3, so-called *fish oils* but algal in origin. Exploitation of the dominant pool of n6 ingested from food to raise the fraction of bio-available n3 counterpart species – lowering the n6/n3 ratio – could form the basis of a cost effective *functional food* supplement and cost-effective therapy for high risk groups in the population. Considering that both n6 and n3 are essential nutrient fats, not intrinsically synthesized that have to be absorbed *across* the gut as free fatty acids and as components within in lipid molecules, the long-term adjustment of the n6/n3 ratio, converting excess n6 to n3 *within* the gut would have far-reaching world-wide public health benefits.

This work considers a vignette, the conflict between indigenous gut bacterial colony and transgenic bacterial colony presented as a live-culture pro-biotic that provides raised n3 nutritional exposure in the lumen of the gut. The introduced transgenic bacteria contributing to raised n3 expresses additional genes for enzymes involved in the conversion of excess n6. The genetic recombinant modifications are for example fatty acid desaturase (FADS) enzymes that convert n6 and illustrated in Figure 1 as red arrows, labelled respectively as Δ^15,17^D, a desaturase converting linoleic acid (LA, C18:2n6) into alpha-linolenic acid (ALA, C18:3n3) and Δ^14,17^D a desaturase converting arachidonic acid (ARA, C20:4n6) into eicosapentaenoic acid (EPA, C20:5n3) and Δ^16,19^D a desaturase converting n6 DPA (Osbond acid, C22:5n6) to DHA(C22:6n3). In a recent review on polyunsaturated fatty acid (PUFA) biotechnology and desaturases, Lee *et al* [8] summarised reports in the literature that such equivalent n6 transformations are possible by selective ω3Desaturase enzymes. Candidate genes for these enzymes are found in microbes and lower eukaryotes (algae) and would be exploitable for altering the n6/n3 ratio in genetic recombinant bacteria. A discussion on the biochemical contributions and genetic re-engineering of ω3Desaturases in addition to the details described in Figure 1, is beyond the scope of this article.

**Figure 1.**
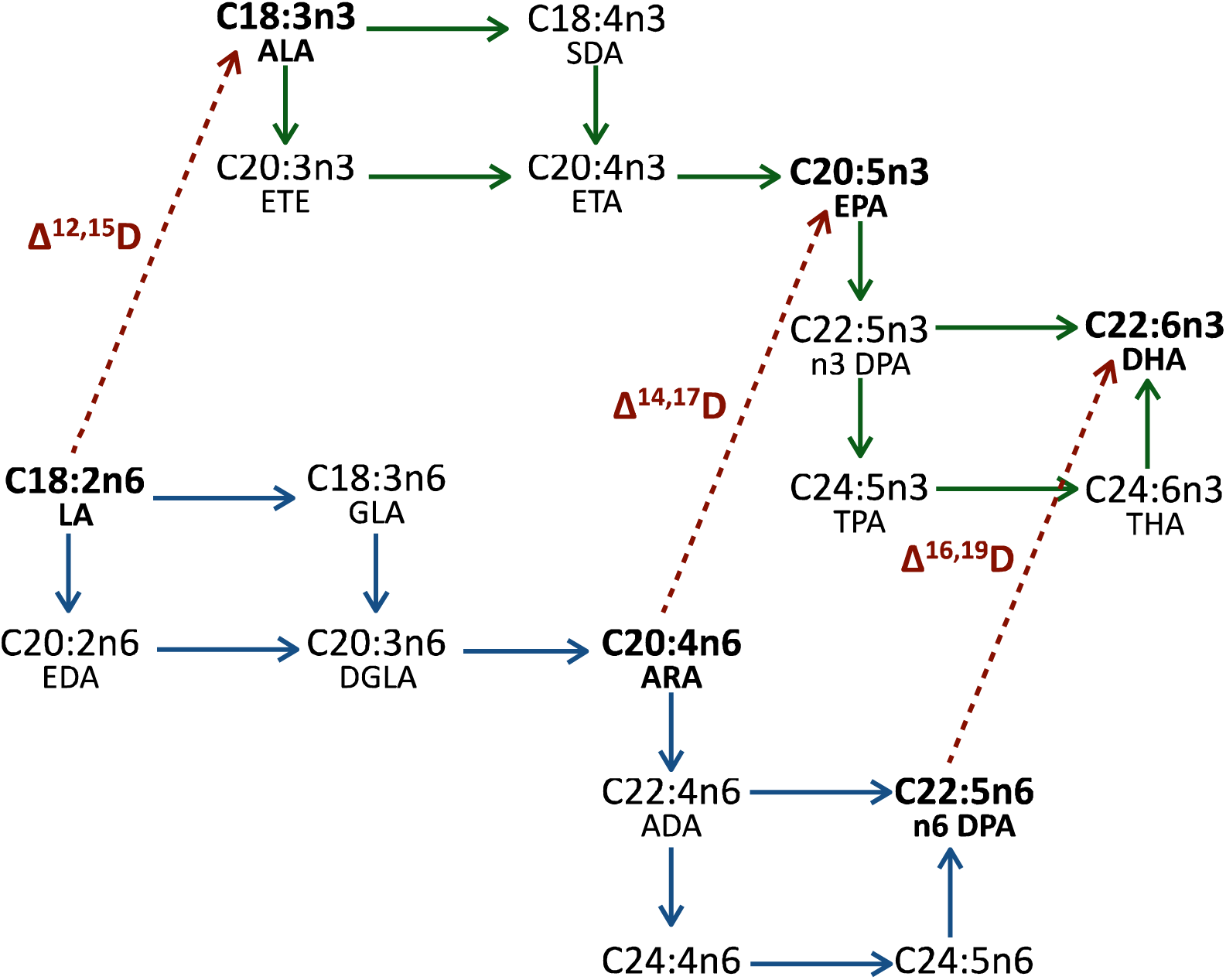
Proposed transgenic ω3Desaturase enzyme transformations (dotted arrows in red) as the potential pro-biotic prophylactic therapy to change the naturally high n6/n3 ratio of polyunsaturated fats (PUFAs). The ω3Desaturase enzymes would act by lowering the abundance of omega 6 PUFAs (n6, network in blue) and raise the abundance of omega 3 PUFAs (n3 network in green). The PUFAs are labelled here by their canonical numbering and labelling and by their abbreviations. The directed networks show the endogenous conversions of the PUFAs and include elongation (down arrows), beta-oxidation (up arrows) and desaturation (left to right arrows), assumed to exist in indigenous gut flora and fauna and in human physiology (further enzymatic details not shown).

Importantly however, we generalise and assume here that the additional genes for the enzyme transformations (illustrated in red) are engineered and collectively provide a transgenic bacterial strain with a new metabolic phenotype with higher abundance of n3 fat-containing components compared to wild type strain. In our vignette, we consider a live culture of the transgenic bacterial colony introduced into the lumen of the gut as a pro-biotic, a functional food or rectal suppository. The application is analogous to the *Caenorhabditis elegans fat-1* gene in Fat-1 mouse and livestock models [9, 10], the introduction of genetically transformed bacteria would certainly have potential to convert n6 to n3 *in situ*. Candidate transgenic bacteria could be selected from symbiotic *Bifidobacterium* already found in the gut, conveniently transplanted or seeded into the intestine of patients [11, 12].

Recognising that the phenotype is stable, optimising inter-microbial interactions that govern the ecological niches between gut microbiota, would be the fundamental challenge for extending the longevity of genetically modified (gut) microbiota therapy [13].

The aim of this work is, therefore, to explore and discuss how a long-term co-existence between native wild type and transgenic bacteria could proceed. A game theoretical approach is ideal for this discussion because it can contribute to predicting the eventual stable endpoints of the populations and help in identifying the strategy (or set of strategies) that cannot be invaded by rare alternative strategies, called the evolutionarily stable strategy (ESS) [14].

## Theoretical Treatment

In the Snowdrift game (or Chicken game, Hawk-Dove game) there is a mixed evolutionarily stable strategy (ESS), where it is always advantageous to adopt the opposite strategy of an opponent. Snowdrift (SD) is illustrated by a scenario, where two car drivers are trapped on either side of a snowdrift and have the options of staying in the car or removing snow from the drift from each of their sides. Allowing the opponent do all the work is the best (selfish) option, however if the opponent remains in their car it is better to shovel, lest no one gets home. Joint co-operation reduces the work load, ensuring that both can drive home.

Should both players co-operate (C, C) then they both receive the reward (*R*) for co-operation. If both players defect (D, D), they both receive punishment (*P*) for mutual defection. Should one defect while their opponent co-operates (C, D or D, C), then the defector receives the “temptation payoff” *T*, while the co-operator receives the “sucker's payoff”, *S* (Table 1).

**Table 1.** The snowdrift game payoff matrix.

|  | Opponent plays |  |
| --- | --- | --- |
|  | Co-operator C | Defector D |
| Focal player plays Co-operator C | $R$ | $S$ |
| Focal player plays Defector D | $T$ | $P$ |

In SD, co-operation yields a benefit *b* to the co-operator as well as to the opposing player and incurs a cost *c* if the opponent defects, but only a cost *c* / 2 if the opponent co-operates, where *b* > *c* > 0. The defector receives the payoff *b* and mutual defection has no net cost nor net benefit, where, *T* = *b*, *R* = *b* – *c* / 2, *S* = *b - c* and *P* = 0 (see Table 2).

**Table 2.** The fitness payoff structure of the snowdrift game.

|  | Opponent plays<br>Co-operator | Opponent plays<br>Defector |
| --- | --- | --- |
| Focal player plays Co-operator | $b - c / 2$ | $b - c$ |
| Focal player plays Defector | $b$ | $0$ |

This leads to the characteristic payoff ranking, *T* > *R* > *S* > *P*. Therefore, it is always better to play as a co-operator when the opponent defects (the probability of encountering a defection increases when defectors are common). Thus, co-operation here is an ESS strategy and co-operators cannot be invaded (*S* > *P*). However, it is also always better to play as a defector when the opponent co-operates (as the probability of encountering co-operation increases when co-operators are common). Thus, defection here is an ESS strategy and defections cannot be invaded (*T* > *R*). Consequently, defectors and co-operators will co-exist. Hence, for getting long-term coexistence between wild type (indigenous) and transgenic bacteria without either one out-competing the other, the two strains would have to engage in an SD interaction.

In our potential pro-biotic prophylactic therapy, the thermodynamic cost of expressing and producing additional ω3Desaturases could decrease the fitness of the genetically engineered (transgenic) bacteria, resulting in their being out-competed by the indigenous gut bacteria (such as *Bifidobacterium*, *Akkermansia muciniphila*, and *Lactobacillus* [15]). The presence of the new ω3Desaturases, however, could also confer a phenotypic advantage. For example, an increase in the n3 would alter the phospholipid components and functionality of the cell wall and its component LPS also known as lipoglycan endotoxin that locally suppresses neighbouring indigenous gut bacteria. Therefore, if the transgenic bacteria were indeed from strain of *Bifidobacterium* or *Lactobacillus*, there could be an adaptive phenotypic advantage when expressing more n3. Although, there would be an energetic cost in producing the new desaturases, there could also be a growth advantage as n3-containing lipid components become more abundant. The transgenic bacteria would play as a co-operator and the wild-type (indigenous) would be the defector. We assume that the indigenous bacteria will not out-compete the transgenic bacteria as in the case of the PD interaction but instead co-exist in the Snow Drift (SD) interaction. We assume that the producers (transgenic bacteria) can benefit from increased n3-containing lipid components before the indigenous bacteria can and that there is net gain from expressing the additional ω3Desaturases. The interaction would convert to the PD interaction [16] if there is a net (energetic or phenotypic disadvantage) in expressing the additional ω3Desaturases, where 2*b* > *c* > *b* instead of *b* > *c* > 0.

The situation above is similar to the competition between the wild type co-operator cells of yeast and the cheater (defector) strains (see, Gore *et al* [17]). Here, co-operator cells produce invertase an enzyme to hydrolyse disaccharide sugars into single sugar residues then 99% of the these spread to and exploited by neighbouring non-producer cells (the cheaters). Yeast cells are known to co-exist and show SD interaction and not PD interaction because the producers reap the profits before the other surrounding cells [17]. Consequently, the SD interactions ensures a longer-term co-existence between the two strains.

Consider a population of *N* members where the fraction of the co-operators is *x* and the fraction of the defectors is 1– *x*; the parameters *T*, *R*, *S* and *P* in (Table 2) can be reduced to a single parameter if *R* = 1 and *P* = 0. The resulting cost-to-benefit ratio *r* = *c* / (2*b* − *c*) and this yields, *T* = 1 + *r* and *S* = 1 – *r*, where 0 < *r* < 1.0.

The equilibrium fractions, from a well-mixed population of co-operators and defectors can be derived using the replicator equation. Replicator dynamics describe the evolution of the frequencies of strategies in a population (from Hofbaueret *et al* [18]), where

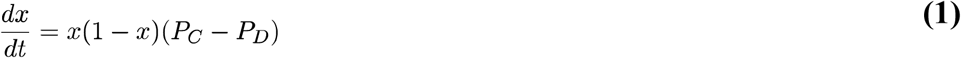

*P*_*C*_ denotes the average payoffs of co-operators,

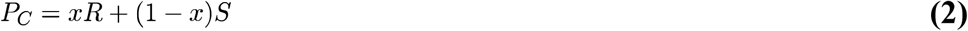

and *P*_*D*_ denotes the average payoffs of defectors,

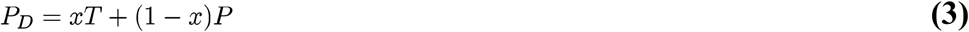

Equation (1) has three equilibrium fixed points *x*_1_ = 0, *x*_2_ = 1 and *x*_3_ where *P*_*C*_ = *P*_*D*_. The third is a stable equilibrium fixed point because the system always returns to it after perturbation, while, the system would move away from the unstable fixed points *x*_*1*_ and *x*_*2*_ after perturbation.

Substituting into Equations (2) and (3) where *R* = 1 and *P* = 0

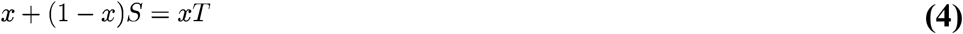

and this leads to

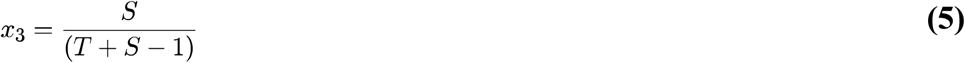

Substituting into the rescaled payoff matrix (Table 3),

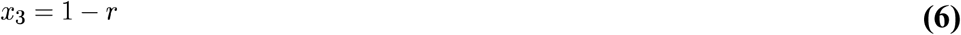

**Table 3.** The rescaled payoff matrix.

|  | Opponent plays<br>Co-operator | Opponent plays<br>Defector |
| --- | --- | --- |
| Focal player plays Co-operator | 1 | 1 - $r$ |
| Focal player plays Defector | 1 + $r$ | 0 |

Therefore, the stable equilibrium of the fraction of co-operators will be

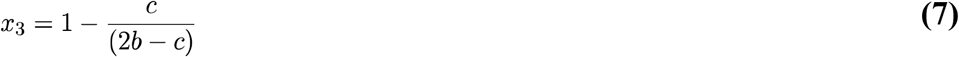

It is interesting that the results of the continuous snowdrift game (CSD) are compatible with the one-shot traditional snowdrift game [19]. The time evolution of the frequency of the co-operators could be shown by numerical simulation at different values of *r*, where 0 < *r* < 1. For instance, if *r* = 0.7, *T* = 1.7, *S* =0.3, *x*_3_ = 0.3. Co-operators and defectors therefore can co-exist in the same population in a stable state of 30% and 70% respectively (Figure 2).

**Figure 2.**
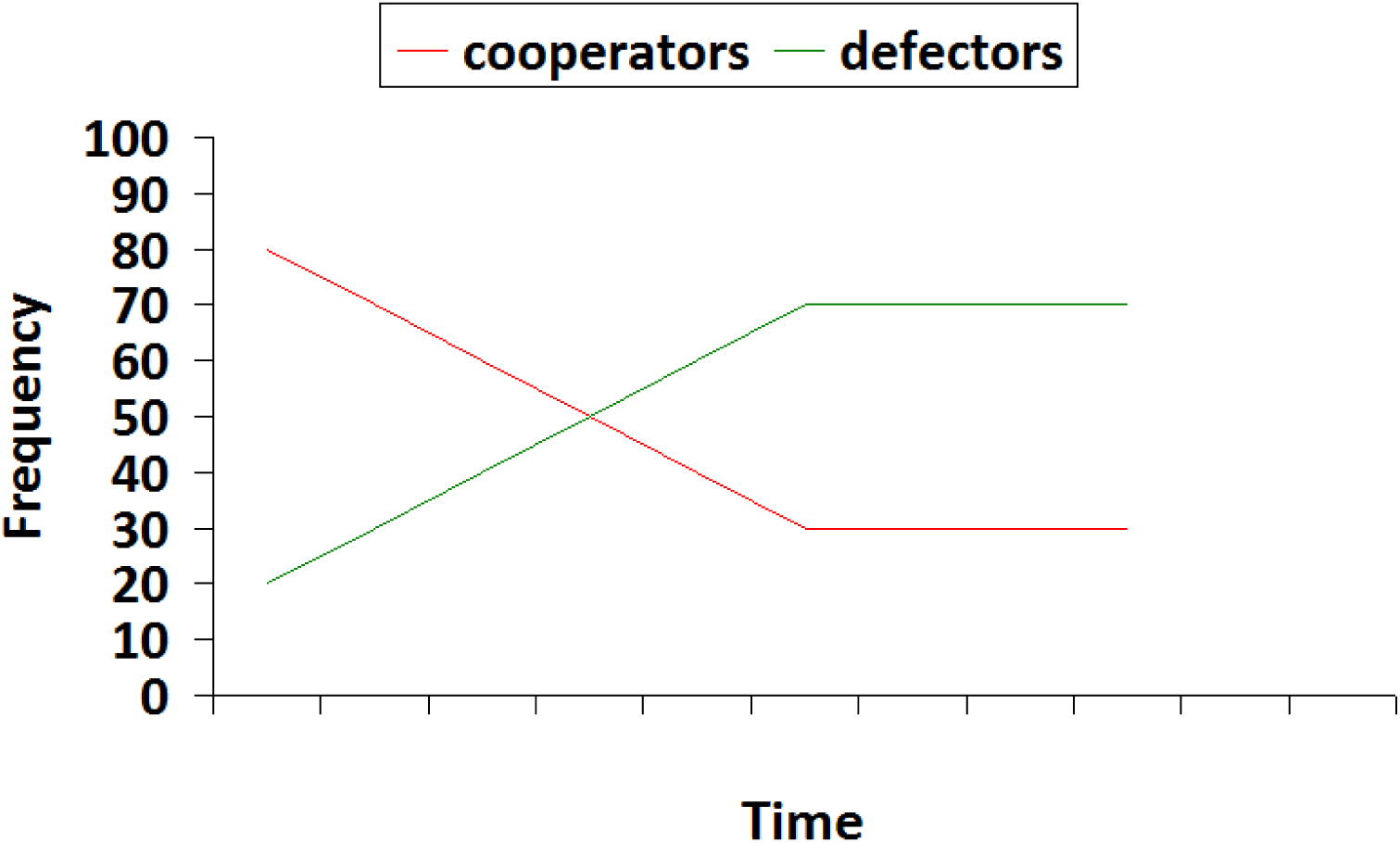
When *r* = 0.7 the equilibrium frequency of the co-operators according to the replicator equation will be 0.30. The stable fraction in the population will be 0.30 co-operation and 0.70 defection, respectively.

**Figure 3.**
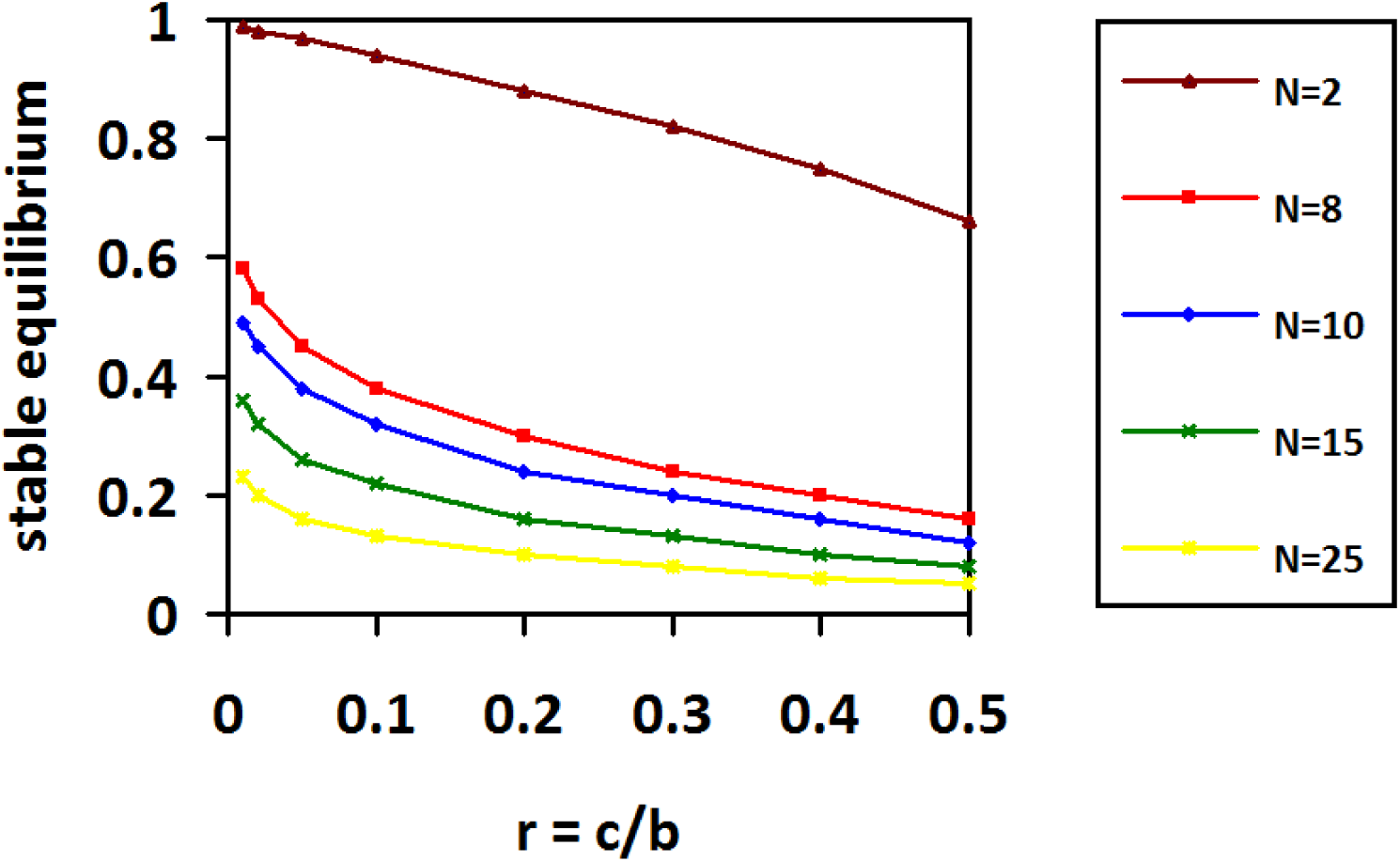
Numerical simulation for Equation (16) illustrating the stable equilibrium of co-operators at different frequencies of low ratios of cost-benefit using random sampling for *N* = 2, 8, 10, 15 and 25.

The interaction (game) will also be PD instead of SD if the co-operation results in a benefit *b* to the opposing player, but incurs a cost *c* to the co-operator (where *b* > *c* > 0), the defector receives the payoff *b*, and mutual defection has no costs or benefits. The payoff rank will be *T* > *R* > *P* > *S*, where *T* = *b*, *R* = *b* – *c*, *P* = 0 and *S* = − *c* (Table 4).

**Table 4.** Fitness payoff of the prisoners’ dilemma.

|  | Opponent plays<br>Co-operator | Opponent plays<br>Defector |
| --- | --- | --- |
| Focal player plays Co-operator | $b - c$ | $-c$ |
| Focal player plays Defector | $b$ | $0$ |

In PD, defection is a pure ESS unbeatable strategy. Therefore, it is always better to adopt the defection strategy regardless of the strategy that the opponent will adopt. Hence defectors will drive co-operators to extinction.

Clearly, there is an empirical problem in determining the payoff structure in social interactions between microbe colonies in nature. Consequently, it is difficult to distinguish experimentally between prisoner's dilemma and snowdrift interactions. Interestingly, work by Fu *et al* [20] suggest an indicator that could solve this problem – Spatial lattice models of macroscopic patterning, exhibiting critical phenomena, reveal that snowdrift interactions have filament-like dendritic clusters, showing percolation behaviour, whereas prisoner's dilemma interactions reveal compact isolated clusters, below a critical propagation threshold.

As an aside, the ability to distinguish emergent patterns of microbial interactions would provide the basis of pattern recognition for colony detection, from faecal microbiology cultures or during endoscopic procedures (for example, with engineered Green Fluorescence Protein in transgenic bacteria [21], borrowing methodological aspects of Borisova *et al* [22] in the clinic, cultured colony detection [23, 24] in the laboratory and colony detection summarised by Hickley *et al* [25]) with which to visualise the spatial-temporal distribution of the most beneficial and effective densities of the probiotic.

After ensuring the outcome of SD, the transgenic bacteria colonies can transmit their effect by providing the omega (n3) fat components to the gut wall. Snowdrift is a two-person game however and realistic interactions are multi-person. We therefore consider Zheng's model of evolutionary snowdrift games with N-person interactions (NESG) [26]. According to the NESG model, in infinite populations without assortments [26, 27] the payoff value for a co-operator is

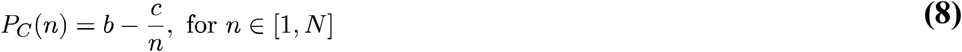

and the payoff value for a defector is

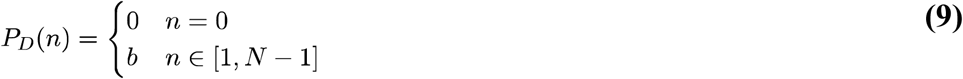

where, *n* is the number of co-operators, represents the number of group members and the benefit (*b*) and cost (*c*) of the collective work. The average fitness of the co-operators and defectors, are respectively,

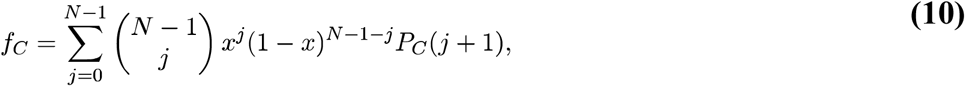

and

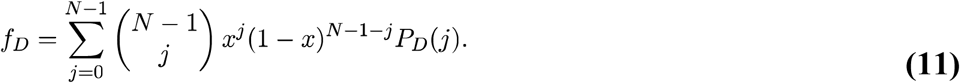

Again, both *x* and (1 - *x*) are the fractions of co-operators and defectors respectively in *N* members but importantly, *j* is the number of co-operators in the group.

The average fitness of the player is

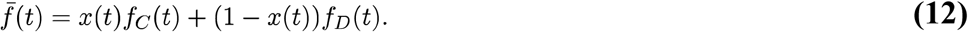

The time evolution of the frequency of co-operation *x*(t) is governed by the following differential equation,

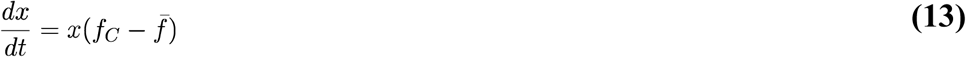

from (1) the replicator equation of frequency of co-operation (t) becomes

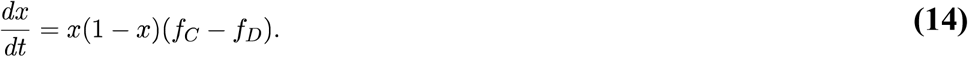

The stable fixed point or the stable equilibrium frequency of co-operation *x*\* satisfies

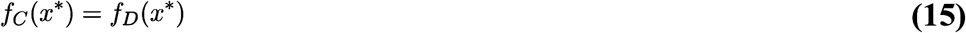

and the stable equilibrium of co-operators can be described by an *N*-th-order equation, Equation (16), where = *c / b* denote to the cost-to-benefit ratio and *x*\* the frequency of co-operation in a group of *N* members.

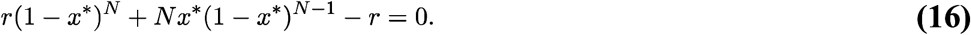

## Discussion

The natural strains of gut microbiota (e.g. *Bifidobacterium*, *Akkermansia muciniphila*, and *Lactobacillus*) have therapeutic effects in a broad range of diseases as probiotics [reviewed in 28, 29, 30] such as inhibiting the growth of harmful bacteria, reducing cholesterol levels, synthesizing vitamins, and treatment of metabolic disorders [31]. Therefore, intake of the genetically modified gut micrbiota as probiotics is considered *per se* a therapeutic mechanism for metabolic syndrome. This feature will not be lost in our model; rather it is supported in an environment rich in n3. Therefore, modified strains expressing ω3desaturases provide a model with multiple therapeutic mechanisms (Figure 4).

**Figure 4.**
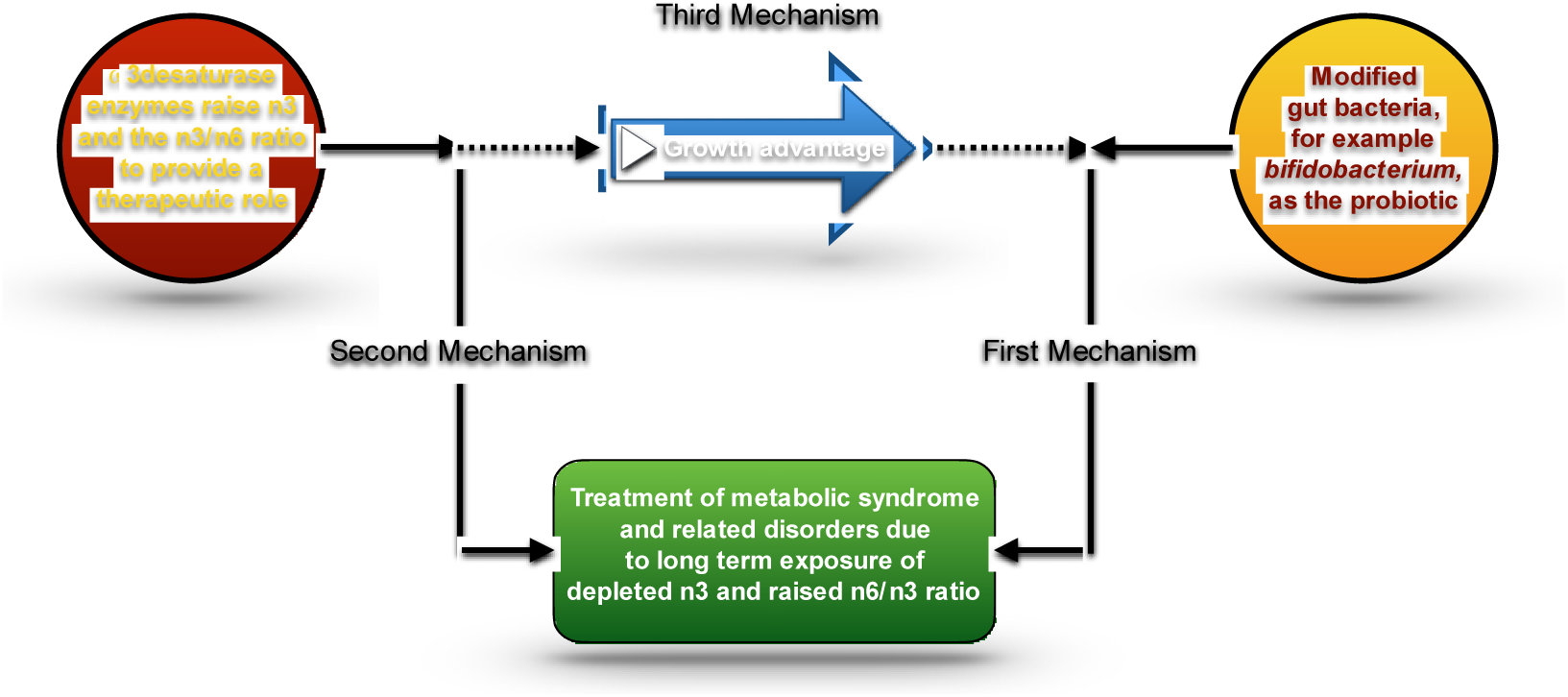
In this model, there are three synergistic mechanisms for treatment of metabolic syndrome and related disorders.

In general, the gram-positive bacteria may have some obstacles to secrete enzymes. Some studies have shown that some strains such as the non-motile *Bifidobacterium longum* can perform this task [32]. Other studies have implicated *Bifidobacterium pseudocatenulatum CECT 7765* strain with a significant therapeutic effect of reducing inflammation and metabolic endotoxemia [33]. Different *Bifidobacterium* species and strains may have different advantages. Hence, we should assume there is a trade-off between all appropriate strains.

### 3.1 Limitations

We used the traditional models of SD and NSD in infinite unstructured populations to shed the light on the possibility of long-term co-existence between the wild type and the transgenic bacteria that in turn are appropriate for prophylactic therapy.

The model does not consider the dynamic of the whole gut species but focuses only on the dynamic between the indigenous and transgenic strain. The assumption in the model is that there are just two strategies interacting with each other, is oversimplified and neglects the contribution of three strategy games [34] (e.g. Rock-Scissors-Paper game) where there are individuals that neither reap benefit nor incur cost. Furthermore, the model does not identify the thresholds of co-operators in the N-person snowdrift game, namely the minimum number of coordinated co-operators required to build the public good [27] and the higher threshold of co-operators beyond which additional co-operators produces no more net benefit [34]. The model also does not consider the evolutionary dynamics of N-person snowdrift game in both structured and finite populations [35, 36].

Although the snowdrift game will lead to co-existence between co-operators and defectors, the spatial structure may suppress the co-operation if it is not dynamic, contributing to the mixing gut microbiota populations [37, 38]. Therefore, co-operators would not be able to develop compact clusters because such patterns increase the proportion of interacting with only other co-operators. Hence, it becomes better to be a defector. Thereby, defectors penetrate co-operators patches and co-operators may extinct if adopt limited dispersal dynamics.

Many theoretical and empirical studies ensure that [39, 40, 41] plasmids which carry cooperative trait can promote co-operation with-in spatial structured patches. In particular, in environmental conditions such as gastrointestinal where the abundance of nutrients [42], by increasing relatedness and homogeneity of the patches via infect the defectors invader with the cooperative traits in the absent of selection for these cooperative traits, provided the null plasmid which carries defection traits was absent or does not have a higher rate of transfer. Consequently, if the rate of infection of defector-invaders by a cooperative mobile element was greater than the rate of colonization of co-operator patches by defectors, the harmful effect of spatial structure on SD game will be averted. Thus, for exceeding the problem of spatial structure we suggest that the cooperative trait (the genes for the desaturases) should be loaded on plasmid.

### 3.2 Future work

From an evolutionary ecological perspective, there are many promising therapeutic applications that can be achieved (for examples, see [43, 44, 45, 46, 47]). If by gaining more insight on heterogeneous microbial interactions and relating them to more sophisticated models of PD and SD-based interactions, critical traits can potentially be eliminated or added to selectively target different patient cohorts.

Subsequently, manipulate the fate of the pathogenic agents leading it to extinction by engaging it in Prisoner’s dilemma [47,48,61] or acquire novel benefits by engaging the engineered beneficial agents in snodrift as in our present work.

All above models address the unconditional deterministic co-operation\defection strategies rather than conditional probabilistic memory-one strategies (e.g. Tit-for-tat (TFT), Win-stay lose-shift (WSLS) *etc.*). A recent revolutionary discovery by Press and Dyson [49, 50] is the conditional zero-determinant (ZD) strategies that allow a player to unilaterally pin the opponent payoff at a certain value or dictate a linear relationship between the own payoff and the opponent payoff, irrespective of the opponent decisions and irrespective of the group size in multiplayer games. ZD strategies are significant powerful long-term control strategies. Interestingly, these strategies are not confined to the iterated prisoner's dilemma or two player games but extended to all public goods games including symmetric social dilemmas [51, 53]. ZD strategies may be fair, extortion or generous.

Individuals adopting extortion ZD strategy reap an unfair share of the payoff from the opponent, on the other hand, they provide incentives to co-operate. Therefore, All-C (the unconditional deterministic strategy that always co-operates) is always the best response to an extortion strategy. Although ZD extortion strategies can dominate any co-operator evolutionary (non-sentient player who accede to extortioner’s actions for maximizing his own score without trying to alter the extortion behaviour) opponent in head-to-head matches, it is still unstable evolutionary strategy as it doesn't act against itself due to mutual defection [53,54]. Therefore, it will win each battle but lose the whole war [55]. For turning it into a stable strategy, extortioners may need for additional mechanisms which enhance the outcomes of interactions between them such as impose players co-operate with themselves but not against the opponent as in coordination games which in turn requires a minimum amount of coordinators. This may be achieved by *i*) tag-based strategies [53] in which tags provide extra information for cooperating only with other extortioners. *ii*) If the extortioners exist in one species which has the slower evolutionary rate of two different interacting species [54] (for example hosts and their symbionts) or two asymmetric classes of the same species.

Generous ZD strategies [56, 57, 58] (compliant strategies) co-operate with others to reap lower payoff than its co-operator co-player and forgive defectors or punish them mildly. A generous ZD strategy is always the best response to itself. Because the baseline payoff (the payoff that a player would get against himself) of compilers is the maximum possible payoff could be reaped in the population (*R*) (no defecting strategy can yield a payoff higher than R when playing against a complier except in the presence of altruistic strategies) [58]. Generous ZD strategies can dominate any defector strategy in the long term even All-D (the unconditional deterministic strategy that always defect). However, unlikely extortioners generous ZD strategies are evolutionarily robust strategy (ERS) [59] and can replace all other classic IPD strategies (e.g. TFT, WSLS, All-D… *etc.*) and even extortion ZD strategies except in very small populations [60].

This is clearly could be a helpful tool for establishing more powerful co-operator transgenic organisms using for the same therapeutic purpose which we suggesting in this article. In general, selection disfavour zero-determinant strategists in large populations [57]. Apparently ZD strategies play an important transient role in the population dynamics, it can be shown in the iterated prisoner's dilemma as the following cycle: both unconditional strategies All-C and All-D enforce linear relationship with a negative slope then the baseline payoff of the whole population decreases over time. Therefore, the population converts from co-operation to defection but extortioners can exist in All-D population by neutral drift then the population converts from defection to co-operation because both conditional strategies extortion ZD and generous ZD enforce linear relationship with a positive slope then the baseline payoff of the whole population increases over time. Eventually, altruistic strategies can invade generous ZD then All-D will rise again (in iterated snowdrift game the slope of All D is positive but the dynamics remain the same) [60].

If tag-based strategies or any other similar strategies succeed to convert extortion ZD to a stable strategy by altering the behaviour of the extortioners against themselves, the extortion ZD will not be anymore the extreme bottom edge of the baseline payoff of the whole population, also it will not be a memory-one strategy. Therefore, it could be used for a therapeutic purpose by outcompeting all of the other strategies, for instance, it can homogenize the colonies of cancer cells or viral strains in the patient. Therefore, it makes them an easy target for drugs or even further it may be invaded by transgenic All-D which imitating the tag [53], then the pathogenic population will be very week or even go extinct (This suggestion is the inverse pattern of our present article).

